# Assessing Bayesian Phylogenetic Information Content of Morphological Data Using Knowledge from Anatomy Ontologies

**DOI:** 10.1101/2022.01.06.475250

**Authors:** Diego S. Porto, Wasila M. Dahdul, Hilmar Lapp, James P. Balhoff, Todd J. Vision, Paula M. Mabee, Josef Uyeda

## Abstract

Morphology remains a primary source of phylogenetic information for many groups of organisms, and the only one for most fossil taxa. Organismal anatomy is not a collection of randomly assembled and independent ‘parts’, but instead a set of dependent and hierarchically nested entities resulting from ontogeny and phylogeny. How do we make sense of these dependent and at times redundant characters? One promising approach is using ontologies—structured controlled vocabularies that summarize knowledge about different properties of anatomical entities, including developmental and structural dependencies. Here we assess whether the proximity of ontology-annotated characters within an ontology predicts evolutionary patterns. To do so, we measure phylogenetic information across characters and evaluate if it is hierarchically structured by ontological knowledge—in much the same way as phylogeny structures across-species diversity. We implement an approach to evaluate the Bayesian phylogenetic information (BPI) content and phylogenetic dissonance among ontology-annotated anatomical data subsets. We applied this to datasets representing two disparate animal groups: bees (Hexapoda: Hymenoptera: Apoidea, 209 chars) and characiform fishes (Actinopterygii: Ostariophysi: Characiformes, 463 chars). For bees, we find that BPI is not substantially structured by anatomy since dissonance is often high among morphologically related anatomical entities. For fishes, we find substantial information for two clusters of anatomical entities instantiating concepts from the jaws and branchial arch bones, but among-subset information decreases and dissonance increases substantially moving to higher level subsets in the ontology. We further applied our approach to address particular evolutionary hypotheses with an example of morphological evolution in miniature fishes. While we show that ontology does indeed structure phylogenetic information, additional relationships and processes, such as convergence, likely play a substantial role in explaining BPI and dissonance, and merit future investigation. Our work demonstrates how complex morphological datasets can be interrogated with ontologies by allowing one to access how information is spread hierarchically across anatomical concepts, how congruent this information is, and what sorts of processes may structure it: phylogeny, development, or convergence.

---

Phylogeny is the key to making sense of biodiversity. It structures the vast variation of form among species into an understandable map that we can use to place and organize all life, compare and contrast organisms, and recover the individual and shared evolutionary history for each lineage and group. By structuring knowledge about data in meaningful ways, a phylogeny allows us to extract information from biological data and ultimately, biological meaning, in ways that would be impossible without it. The hierarchical nature of life, however, is evident not just at the level of species (e.g., Oakley 2003; Serb and Oakley 2005). It is also observed among phenotypic traits, which are themselves often descended from common ancestral precursors modified over developmental and evolutionary time frames.

Therefore, organismal anatomy is not a collection of randomly assembled ‘parts’. It is the manifestation of relationships among anatomical entities and structure resulting from ontogeny and phylogeny. Just as we can organize knowledge about species with phylogeny, our definitions of the entities, qualities, and relations of organismal traits can be organized by ontologies—structured controlled vocabularies formalizing relationships among concepts (Mabee et al. 2007; Vogt 2009; Deans et al. 2015).

Ontologies summarize knowledge about different properties of anatomical entities, including developmental and structural dependencies. For example, in fishes, the presence of a ‘dorsal fin ray’ is dependent on the presence of a ‘dorsal fin’. Here, we explore one particular aspect from ontologies: do ontology concepts referring to real anatomical entities and the relations among them structure phylogenetic information? In other words, does the proximity of characters within an ontology with respect to their anatomical and structural relations predict their evolutionary patterns? Investigating this question is key to understanding the processes underlying morphological evolution and to addressing key impediments to the ‘Phenomics’ revolution (Deans et al. 2015)—namely, the complex sets of dependencies among phenotypic characters that confound the application of traditional statistical models.

In contrast to molecular data, which are typically treated as independently-evolving sets of characters, morphological data are known to carry dependencies and redundancies across characters. Morphological traits may change in a concerted fashion through evolutionary time (i.e., evolutionary modules) if they share a common underlying genetic/developmental machinery (Lewontin 1978; Wagner 1989, 1996, 2007; Wagner and Altenberg 1996; Wagner and Stadler 2003; Mabee 2006) and/or as a result of shared functional/ecological selective pressures (e.g., see concerted convergence: Patterson and Givnish 2002; Holland et al. 2010; Blank et al. 2013). Therefore, groups of characters may imply similar trees due to shared phylogenetic history (Fig. 1a) or convergence (Fig. 1b), and in both cases may over-represent the degree of support if treated as independent realizations of a stochastic evolutionary process. In this context, ontology knowledge may provide us with additional insights (e.g., from anatomy and development) into the historical patterns of trait changes: Do particular classes of anatomical entities provide more phylogenetic information than others (Fig. 1c)? How semantically diverse are the anatomical concepts that support a particular topology? Is there conflict between different sets of anatomical concepts that may suggest convergence or other evolutionary processes (Fig. 1d)?

**FIGURE 1.**
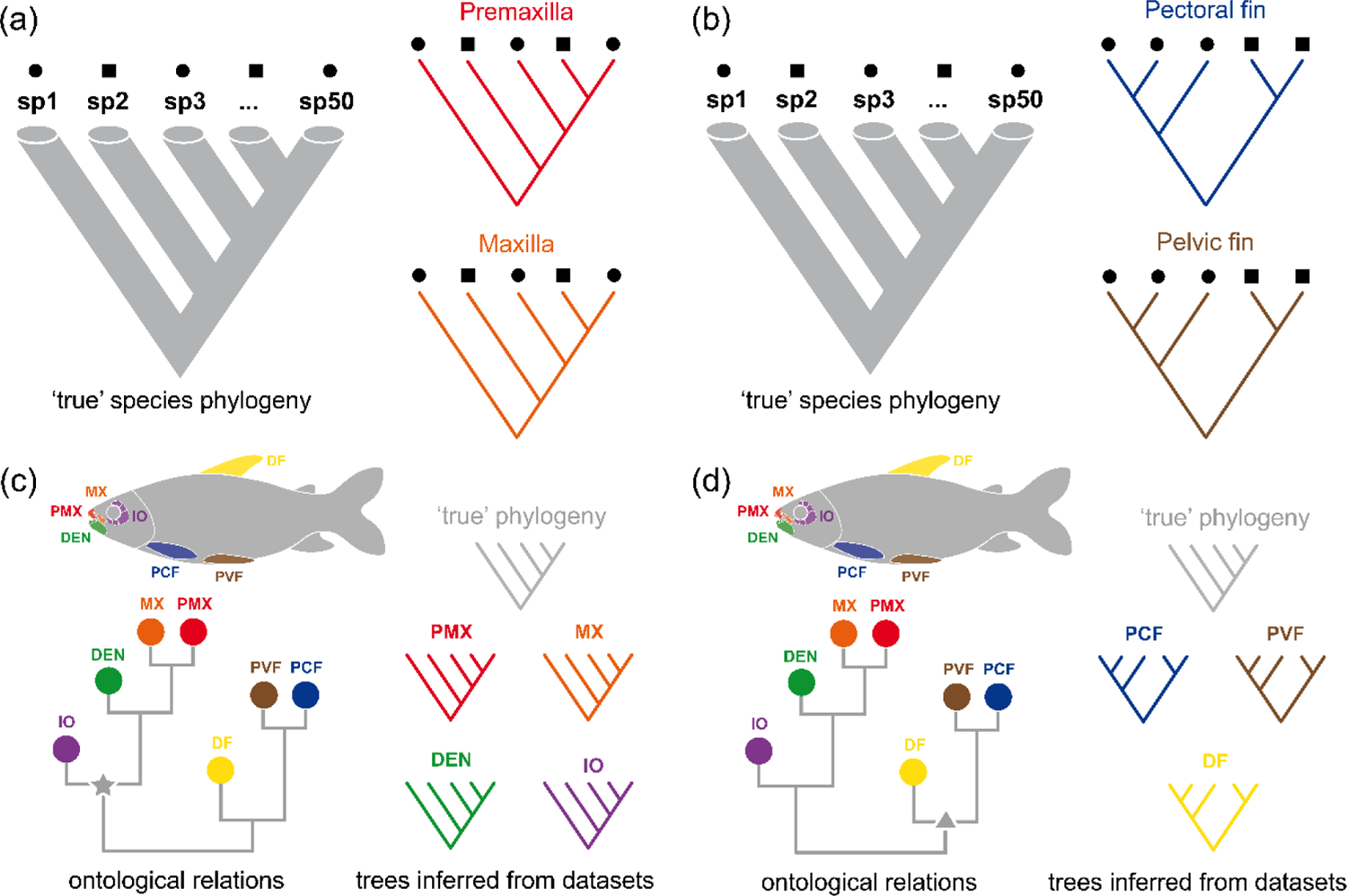
Comparison of the ‘true’ species phylogeny with trees inferred from different data subsets. (a) Trees inferred from characters of ‘premaxilla’ and ‘maxilla’ are congruent and indicate true phylogenetic information. (b) Trees inferred from characters of ‘pectoral fin’ and ‘pelvic fin’ are congruent between themselves but not with the ‘true’ species phylogeny, thus indicating convergence, in this case, associated with other ecological/functional factors (squares and circles). (c) Ontology relations among anatomy entity concepts showing that related anatomical entities (for example, the node indicated with a star) provide true phylogenetic information. (d) Ontology relations among anatomy entity concepts showing that related anatomical entities (for example, the node indicated with a triangle) provide no phylogenetic information, but are jointly influenced by convergent evolution. Abbreviations: DF, dorsal fin; DEN, dentary; IO, infraorbital; MX, maxilla; PCF, pectoral fin; PMX, premaxilla; PVF, pelvic fin; sp1…sp50, species in a dataset. For colors, please refer to the online version of this paper available at XXX.

Here, we develop a view that is distinct from typical partitioning of phylogenetic datasets. Approaches to assess and/or account for heterogeneity across subsets/partitions of molecular data (e.g., genes, codon positions) usually focus on rates and/or model of trait evolution (see review in Kainer and Lanfear 2015); informativeness (e.g., Townsend 2007; Townsend et al. 2012); or topological conflict among inferred trees (e.g., Zhou et al. 2020; Smith et al. 2020). However, much like how partitioning taxa into a flat set of genera or families is inadequate to represent phylogenetic structure, partitions in the traditional sense fail to account for the continuous hierarchical relations among characters. Expanding partitions among characters into hierarchical structures enables new questions to be asked of phylogenetic data.

One such fundamental question is to ask how information about the phylogeny is structured across characters. Here, we address this question by integrating knowledge from ontologies with the Bayesian phylogenetic information (BPI) framework proposed by Lewis et al. (2016) (see also Neupane et al. 2019; Porto et al. 2021). Lewis’ et al. framework is based on Shannon’s (1948) entropy and Lindley’s (1956) information. In short, Shannon’s entropy measures uncertainty in discrete outcomes and Lindley’s information measures how data make some outcomes more probable than others. In Lewis’ et al. context, the outcomes refer to discrete tree topologies in the posterior distribution. Therefore, (Bayesian) phylogenetic information is used here in a sense that differs from most common usages.

Phylogenetic information usually refers to the information inferred from data about the ‘true’ evolutionary history of organisms (see discussion on phylogenetic systems in Farris 1979). A related concept is phylogenetic signal, which refers to similarity among an organismal trait (or set of traits) in different taxa that is explained by shared evolutionary history (Pagel 1999). BPI here refers to the ‘ability’ of data to concentrate prior probabilities of tree topologies into a smaller set of trees in the posterior (as in Lewis et al. 2016).

Lewis’s et al. approach allows us to assess information inferred from data, but also to evaluate how different data subsets may concentrate probabilities into alternative sets of trees through a measure called phylogenetic dissonance (Lewis et a. 2016). Data subsets may represent groups of characters from different anatomical regions. They can be compared to evaluate which ones are congruent with each other and/or with the ‘true’ phylogeny, for example. Ontology knowledge can be integrated by structuring such comparisons in a meaningful way based on known relations (e.g., anatomical/developmental) among anatomy entity concepts instantiated by characters annotated in these data subsets (Fig. 2a). Semantic similarity then can be employed to assess how closely related two anatomical concepts are in the ontology, a metric that can be used to link characters in a character matrix to a ontologically structured hierarchy, thus providing the backbone for comparisons among data subsets (Fig. 2a).

**FIGURE 2.**
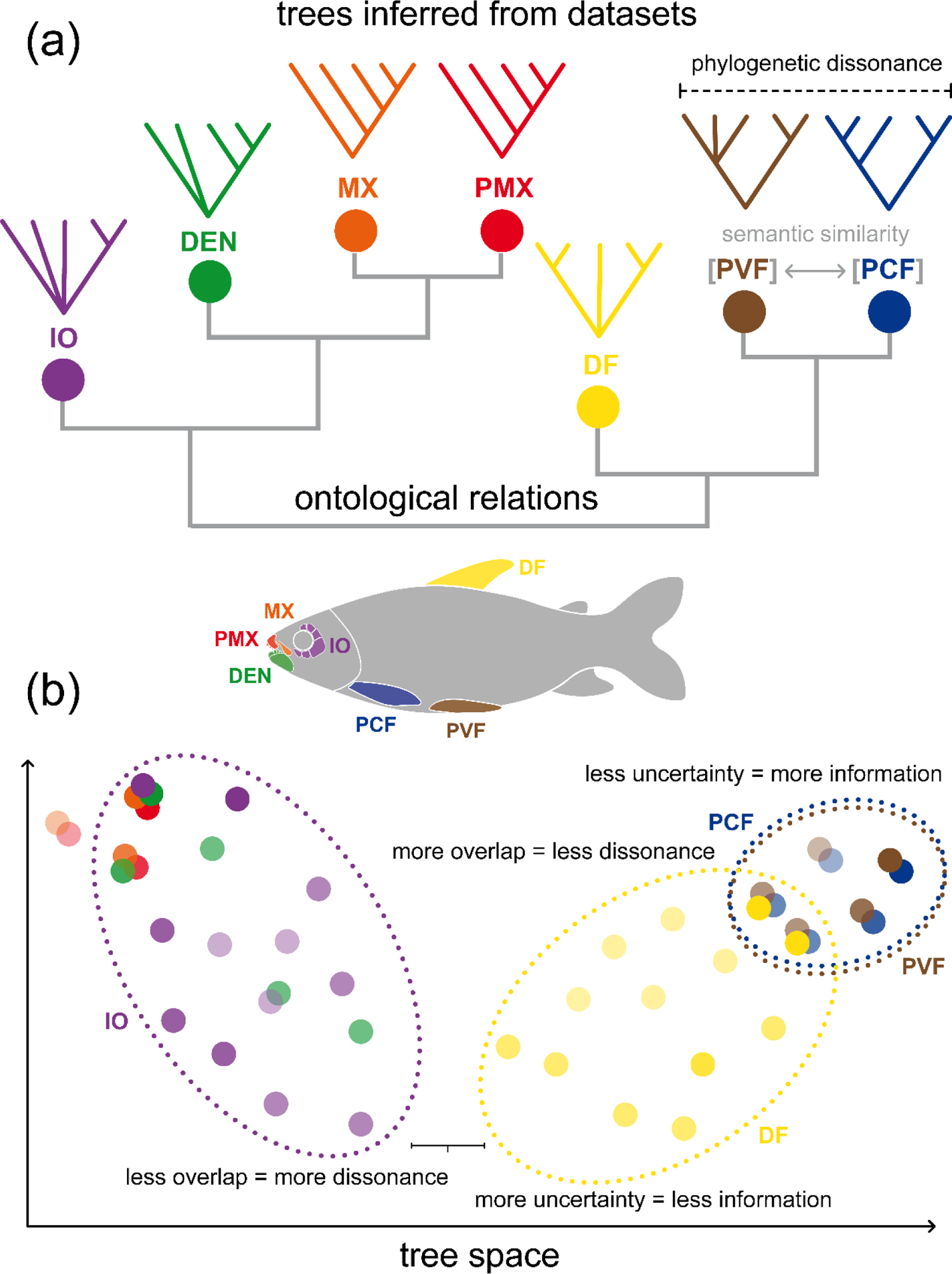
Diagrammatic representation of the relationship between ontology structure, represented as a clustering dendrogram, and a hypothetical posterior tree space. (a) Ontology hierarchy of anatomy entity concepts referring to data subsets used to infer posterior distribution of trees (only one tree shown above each term). The hierarchy is represented as a clustering dendrogram based on semantic similarity distances among anatomy entity concepts (b) Representation of a hypothetical posterior tree space. Each circle indicate a discrete tree topology. Shade intensity is proportional to the posterior probability of each topology. Dotted ellipses indicate the hypothetical area of the tree space occupied by inferred trees in the posterior of some data subsets. Abbreviations: DF, dorsal fin; DEN, dentary; IO, infraorbital; MX, maxilla; PCF, pectoral fin; PMX, premaxilla; PVF, pelvic fin. For colors, please refer to the online version of this paper available at XXX.

The approach advocated here combines elements of information theory with ontology knowledge allowing one to investigate what sort of processes may structure probabilities in the tree space of the posterior distribution of tree topologies (Fig. 2b). BPI provides a measure of how much uncertainty there is in the posterior inferred from a data subset: lower BPI means more possible trees with probability scattered across them (e.g., Fig. 2b: DF); higher BPI means fewer possible trees with probability concentrated in some of them (e.g., Fig. 2b: PVF or PCF). Phylogenetic dissonance provides a measure of how congruent posterior distributions of trees inferred from different data subsets are: lower dissonance means that a similar set of trees with similar probabilities are present in the posteriors (e.g., Fig. 2b: PVF vs. PCF); higher dissonance means that there is low or no overlap among the posteriors (e.g., Fig. 2b: DF vs. IO). If subsets are defined based on organismal anatomy and the patterns of phylogenetic dissonance observed in the tree space of the posterior (Fig. 2b) reflect the ontological hierarchy (Fig. 2a), then one can ask if anatomy/development may play a role in explaining phylogenetic information in the data. In other words, ontology knowledge structures phylogenetic information in this case (i.e., there is *semantic signal*). Alternatively, if groups of unrelated anatomy entity concepts (i.e., low semantic similarity) provide congruent trees and such trees are congruent with the ‘true’ species phylogeny (Fig. 1a), then such entities are just following the common species history. Finally, if groups of unrelated anatomy entity concepts provide congruent trees that are different from the species tree (Fig. 1b), then other processes may be suspected (e.g., concerted convergence).

In this study we evaluated the phylogenetic information content of ontology-annotated character matrices by measuring BPI and phylogenetic dissonance applied to morphological data in phylogenetic inferences. We applied this approach to two datasets representing disparate animal groups for which well-established anatomy ontologies are available: bees (Hexapoda: Hymenoptera: Apoidea) and characiform fishes (Actinopterygii: Ostariophysi: Characiformes). Within the characiform fishes, we further targeted specific evolutionary questions concerning miniaturization, which is predicted to result in convergent evolution among certain data subsets. We propose a new framework for evaluating alternative hypotheses for the sets of ontological relationships that best explain phylogenetic information across ontology-annotated anatomical data subsets (i.e., *semantic signal*). This framework is not limited to the Bayesian information metrics used here, and it can be a general approach to understanding how ontologies may structure phylogenetic information inferred from anatomical data and investigating whether morphologically related entities show similar tree-like histories due to a shared phylogeny (i.e., *phylogenetic signal*) or other process such as concerted convergence. We have made our implementation of this methodology available in the new R package *ontobayes* (https://github.com/diegosasso/ontobayes).

## MATERIAL AND METHODS

### Theoretical Background

#### Definitions

Throughout this paper, we employed a few terms with varying usage in the literature. ‘Dependency’ (e.g., either anatomical, morphological, or structural) is used in the same sense as ‘ontological dependency’ (Vogt 2018a) to describe the types of relationships when the absence/presence of one anatomical entity determines the absence/presence or condition of another. The terms ‘trait’ and ‘character’ are used mostly interchangeably to mean “any recognizable phenotypic unit from organisms”. Here, ‘character’ is used to specifically refer to phenotypic units that are variable across organisms and used as input data in phylogenetic analyses. We make a distinction in the use of the terms ‘dendrogram’ and ‘tree’, despite the former including the latter. ‘Dendrogram’ or ‘clustering dendrogram’ is used here to refer to any tree-like hierarchical diagram depicting relationships among anatomy ontology terms. ‘Tree’ or ‘phylogenetic tree’ is reserved to the hierarchical diagrams depicting relationships among species. ‘Topology’ is used to refer to the ordering of the hierarchy among leaves in such tree-like diagrams, without respect to edge length. ‘Term’ is used to refer to the labels applied to real anatomical entities represented as concepts in an anatomy ontology. ‘Data subset’ or ‘partition’ is used to refer to groups of traits/characters annotated with or descended from a particular ontology term/concept.

#### Ontologies

Ontologies are structured controlled vocabularies formalizing relationships among concepts in a specific domain of knowledge, for example, vertebrate (Dahdul et al. 2012; Haendel et al. 2014) and hymenopteran anatomy (Yoder et al. 2010). Concepts can be expressed by terms linked to or defining organismal anatomical entities (e.g., ‘opercle’ from the Uberon anatomy ontology, Mungall et al. 2012; Haendel et al. 2014) or phenotypic qualities (e.g., ‘triangular’ from the Phenotype and Trait Ontology, Gkoutos et al. 2005), and phenotypes can be described using the Entity-Quality syntax (e.g., E: ‘opercle’, Q: ‘triangular’) (Mungall et al. 2010; Balhoff et al. 2010; Dahdul et al. 2010a, 2012). Relationships among concepts can be of various kinds (e.g., *part_of*, *is_a*, *develops_from*) and different logical relations may be included to build knowledge graphs with relevant structural or developmental information about organismal traits (e.g., Dahdul et al. 2010b; Mabee et al. 2012).

Ontological knowledge can be explored in different ways to summarize information on structural dependencies among anatomical entities instantiating ontology concepts. One possibility is to use semantic similarity measures to build a dendrogram depicting distances among anatomy entity concepts (Fig. 2a). Semantic similarity can be assessed using different metrics such as edge-based distances (e.g., *Jaccard*), node-based information content (e.g., *Resnik*), or hybrid metrics (e.g., Hybrid Relative Specificity Similarity)(Pesquita et al. 2009; Manda and Vision 2018). Different metrics can capture alternative and/or complementary properties of the ontology. The types of relations included as well as the ontology structure itself can influence the overall similarity values between concepts (Pesquita et al. 2009; Manda and Vision 2018). Another possibility is to use ontological knowledge to explicitly account for anatomical dependencies among individual traits when specifying models of character evolution (Tarasov 2019, 2020; Tarasov et al. 2019). This can be achieved by constructing models of discrete trait evolution enabling ontology-aware transition matrices through structured Markov models equipped with hidden states (Tarasov 2019). In this work, we focused on the first way of exploring ontology knowledge.

#### Bayesian phylogenetic information

BPI is the amount of information about phylogenetic tree topology inferred from the data. It is measured as the difference in entropy between prior and posterior probability distributions on phylogenetic tree topologies (Lewis et al. 2016). In this context, entropy can be interpreted as a measure of uncertainty and is inversely proportional to information. If data provides no information in favor of any phylogenetic tree topology, then entropy (and uncertainty) is maximal and all possible trees are equiprobable (assuming a discrete uniform prior). Thus, phylogenetic information inferred from data will make some phylogenetic tree topologies from the prior more probable than others resulting in a concentrated posterior (Lewis et al. 2016).

Comparing BPI from different subsets allows the estimation of the amount of informational conflict between posterior probability distributions of phylogenetic tree topologies—i.e., phylogenetic dissonance (Lewis et al. 2016; Neupane et al. 2019). In this study, we asked whether ontology structures Bayesian phylogenetic information for phylogenetic tree topology (see also Lewis et al. 2016; Neupane et al. 2019; Porto et al. 2021), although similar questions could be asked for other types of information (e.g., regarding ancestral states for discrete characters; Borges et al. 2019). Since the prior on phylogenetic tree topology for a dataset with a given number of taxa is the same as for all its possible subsets—for unrooted dichotomous labeled phylogenetic trees it depends only on total number of taxa—BPI from different data subsets can be compared to assess their individual informational contributions in phylogenetic analyses (e.g., Neupane et al. 2019; Porto et al. 2021). Therefore, BPI and phylogenetic dissonance provide straightforward measures for assessing how much agreement or disagreement there is between the posterior distributions of phylogenetic tree topologies from two or more data subsets. We use these statistics to investigate how the ontology structure translates as congruence or dissonance in phylogenetic information provided by different subsets.

#### The Ontobayes Approach

To measure phylogenetic information and dissonance, we carried out four main steps using R (R Core Team 2021), MrBayes (Ronquist et al. 2012), and Galax (Lewis et al. 2016) in an implementation of our analysis, which we call *ontobayes*. In brief, we aggregate ontology-annotated characters into subsets based on anatomical terms and use phylogenetic analyses of these subsets in MrBayes to obtain posterior samples of phylogenetic tree topologies. The samples are then used to calculate the information theory metrics (i.e., BPI and phylogenetic dissonance) in Galax to compare different subsets. All functions, examples, and documentation for the *ontobayes* R package are available at https://github.com/diegosasso/ontobayes and in the online Supplementary Material available on Dryad.

We incorporated ontological knowledge about organismal anatomy (Fig. 3a) by building data subsets (Fig. 3d) grouping characters based on ontology term annotations (Fig. 3b) and structuring relationships among ontology concepts as clustering dendrograms (Fig. 3c). We based dendrograms on distance matrices from measures of semantic similarity using functions from *rphenoscape* (https://github.com/phenoscape/rphenoscape) (Fig. 3c, hereafter ‘semantic similarity dendrogram’) or phylogenetic dissonance (hereafter ‘dissonance dendrogram’). We evaluated two alternative ways of constructing dendrograms based on: (1) all available terms annotated to characters in a given character matrix (ALL) and (2) a smaller selection (‘profile’) of preferred terms (PROFILE), which allow for specific investigation of terms of particular research interest. We estimated BPI and phylogenetic dissonance in Galax (i) among different MCMC runs from the same data subset and (ii) from different data subsets. The former analysis assess the topological convergence and information content (Lewis et al. 2016), while the latter measures concordance or conflict among two or more distinct data subsets. Entropy measures in Galax (Fig. 3e, e.g., E1 and E2) were then be used to estimate information content and conflict by assessing uncertainty in posterior probability distributions (see discussions in Lewis et al. 2016; Neupane et al. 2019; Porto et al. 2021).

**FIGURE 3.**
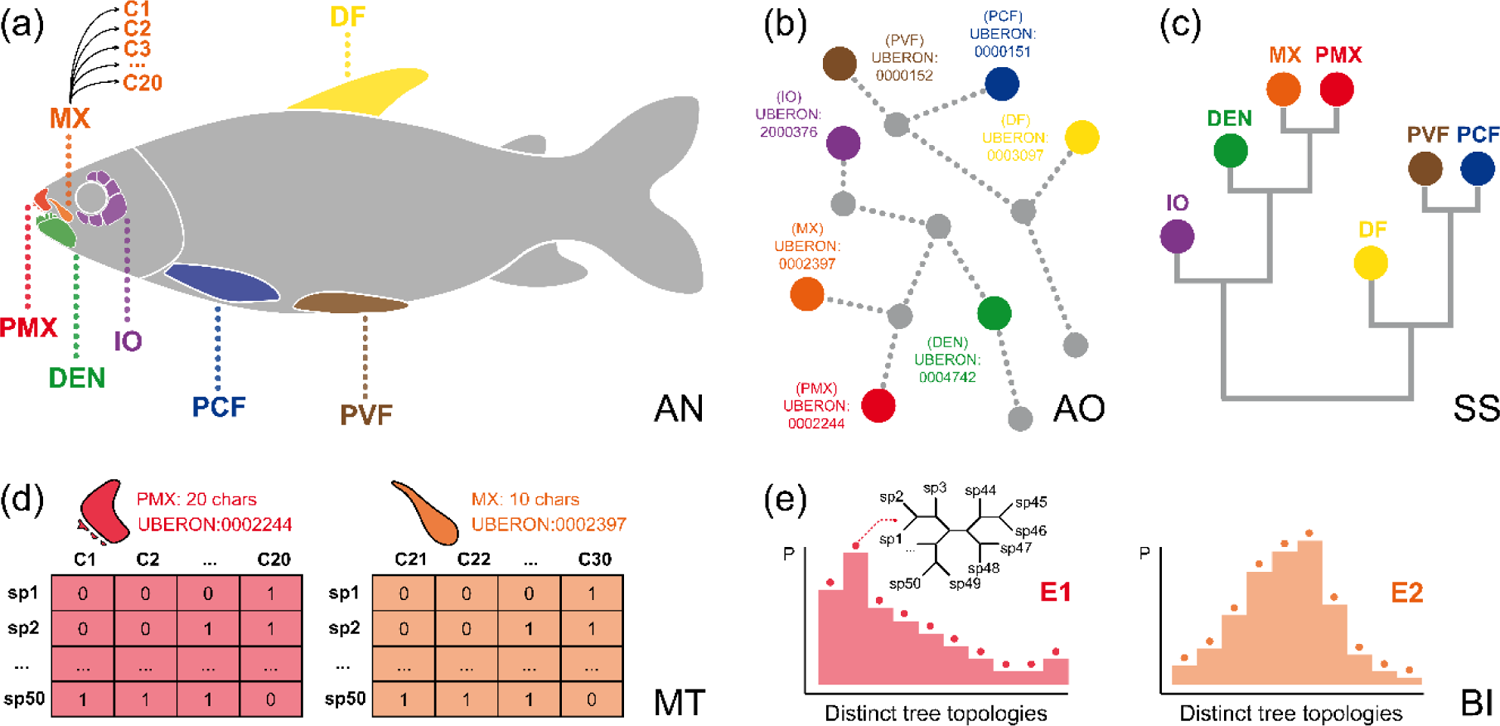
Diagrammatic representation of main steps of the *ontobayes* analysis. (a) Ontology terms referring to anatomical entities are linked to characters in a matrix using expert judgment. (b) Terms in the ontology are related to other terms by logical relations (e.g., *is_a*, *part_of*) which can be represented as a graph. (c) Semantic similarity metrics derived from such a graph (e.g., *Jaccard*, *Resnik*) can be employed to build a clustering dendrogram for terms. (d) The structure of such a dendrogram can then be used to guide comparison of subsets of characters linked to the same or related ontology terms. (e) Each subset is used to produce posterior probability distributions of phylogenetic tree topologies which are used to estimate Information Theory metrics (i.e., entropy, information, dissonance). Abbreviations: AN, organismal anatomy; AO, anatomy ontology; BI, Bayesian inference; C1…C30, characters in a matrix; DF, dorsal fin; DEN, dentary; E1…E2, entropy of posterior distributions; IO, infraorbital; MT, character matrices; MX, maxilla; PCF, pectoral fin; PMX, premaxilla; PVF, pelvic fin; sp1…sp50, species in a matrix; SS, semantic similarity dendrogram. For colors, please refer to the online version of this paper available at XXX.

#### Empirical Analyses

We analyzed how ontology structures phylogenetic information with two datasets representing disparate animal groups: bees and characiform fishes. The two groups were selected for this study since well-established anatomy ontologies are already available for them (bees and other hymenopteran insects: Hymenoptera Anatomy Ontology, HAO, Yoder et al. 2010; vertebrate animals: Uberon anatomy ontology, Mungall et al. 2012; Haendel et al. 2014) and comprehensive character matrices could be annotated with ontology terms based on the authors’ expertise. The BEE dataset was modified from Porto et al. (2021), which includes corbiculate bees and related taxa (Hexapoda: Hymenoptera: Apidae). The original matrix was reduced to contain only 10 bee species because Bayesian phylogenetic information content estimation is less reliable when the number of taxa (and thus possible phylogenetic tree topologies) is large (Lewis et al. 2016). Two species representing each of the four corbiculate bee tribes (i.e., Apini, Bombini, Euglossini, and Meliponini) were selected, plus two outgroup taxa (Centridini: *Epicharis* and Anthophorinae: *Anthophora*). The taxon sampling represents the diversity among the main lineages of Apinae bees (e.g., see Porto et al. 2021). The final dataset contained a total of 209 informative characters, each annotated with anatomical terms from HAO (see supporting data in the online Supplementary Material available in Dryad).

This dataset was first analyzed under the PROFILE alternative of subset construction, which is based on pre-defined groups of selected ontological terms (i.e., ‘profiles’). Six groups of terms from HAO were chosen so as to assess the information content and dissonance within and across data subsets representing groups of anatomical entities in distinct body regions from the bee anatomy. The anatomical terms were selected so as to represent the main morpho-functional regions in the body of a typical apocritran Hymenoptera. The groups of selected terms were: 1. Mouthparts: labrum, mandible, maxilla, labium, and sitophore; 2. Head: cranium and tentorium; 3. Mesosoma: prothorax, mesothorax, and metathorax; 4. Legs: fore, mid, and hind legs; 5. Wings: fore and hind wings; 6. Metasoma: male and female genitalia. In addition to these pre-defined profiles, we analyzed this dataset under the ALL alternative of subset construction to obtain a dissonance dendrogram, which represents relationships among all ontological terms annotated to characters in the matrix by estimating phylogenetic dissonance for all pairwise comparisons among data subsets (see supporting data in the online Supplementary Material available in Dryad).

The FISH dataset was obtained from Dillman et al. (2016) and includes information for four families of anostomoid fishes in the order Characiformes (Actinopterygii: Ostariophysi: Characiformes). The original matrix was reduced so as to contain only 10 taxa and retaining 463 characters, each annotated with anatomical terms from the Uberon ontology (see supporting data in the online Supplementary Material available in Dryad). Two or more species representing the four anostomoid families (i.e., Anostomidae, Chilodontidae, Curimatidae, and Prochilodontidae) were selected, along with one outgroup taxon (Parodontidae: *Parodon*). The taxon sampling represents the diversity among the main lineages of anostomoid fishes (e.g., see Dillman et al. 2016). This dataset was analyzed under the ALL alternative of subset construction. It was first used to compare alternative ways of representing the relationships among subsets of ontological terms (e.g., phylogenetic dissonance and semantic similarity dendrograms) to assess congruence between ontology structure and phylogenetic information. We then evaluated (1) the information content of individual data subsets defined as groups of characters annotated to the same ontological term; (2) information content and dissonance among distinct subsets defined by different ontological terms; and (3) clade-specific information components provided by each subset to nodes in a given reference phylogenetic species tree. The reference species tree was inferred using all 463 characters for the 10 fish species sampled and represents the phylogenetic knowledge acquired when the information from all characters annotated to all ontological terms is considered together. The dataset was also used to investigate whether subsets of data might be ontologically related to particular terms (in this case, as an example, all terms that are *part_of* ‘dermatocranium’).

Within characiform fishes, miniaturization has occurred multiple times and may result in convergent character states for sets of traits. To evaluate whether the degree to which characters respond to these convergent selection pressures is structured by ontology we assembled a modified third dataset, the MINI dataset, from Mirande (2019). We focused on 10 species of characiform fishes that had multiple convergent miniatures and retained 453 characters, each annotated with anatomical terms from the Uberon ontology (see supporting data in the online Supplementary Material available in Dryad). Specifically, taxon selection included: four miniature fishes (body size < 26mm *sensu* Weitzman and Vari 1988) and six non-miniature fishes representing four different lineages of Characidae and two outgroups. Each characid lineage was represented by a miniature and a non-miniature species. To assess convergence, a reference phylogenetic tree was inferred that constrained all miniatures to a monophyletic grouping. Clade-specific information components were then obtained for all subsets of characters to determine whether the ontology structures which traits are more informative about miniaturization phenotypes, and which traits follow the species tree.

Finally, we further evaluated whether the ontology structures phylogenetic information and dissonance across characters by conducting comparisons of ontology-based subsets and randomly resampled sets of characters from the original FISH dataset. Six terms from the Uberon ontology annotated to multiple characters in the FISH dataset and representing different anatomical entities (i.e., fish bones) were chosen: 1. Premaxilla (PMX, 8 chars), 2. Maxilla (MX, 14 chars), 3. Dentary (DEN, 8 chars), 4. Infraorbital (IO, 11 chars), 5. Epibranchial bone (EB, 15 chars), and 6. Ceratobranchial bone (CB, 10 chars). For each term, 100 different subsets of the same size were produced by randomly sampling characters from the original FISH dataset. These resampled subsets were compared to the ontology-based (hereafter ‘standard’) subsets by measuring BPI and phylogenetic dissonance. Ontological relationships among selected terms were represented as a semantic similarity dendrogram that was then employed to guide sequential pairwise comparisons between data subsets based on terms with successive increasing distances within the ontology (i.e., decreasing semantic similarity) adopting one term as a fixed reference (i.e., PMX, premaxilla).

Posterior samples of phylogenetic tree topologies were obtained running MCMC analyses in MrBayes (Ronquist et al. 2012) with two runs and four chains, for 1.0 × 10^7^ generations, sampling every 1000^th^ generation, and discarding the first 25% as burn-in. The Mk+G model was employed with the following priors and parameters: 1. Tree topology prior: Discrete Uniform (1, |T|); 2. Branch lengths prior: Exponential (10); 3. Discrete Gamma shape: Exponential (1); 4. State frequencies: Symmetric Dirichlet (infinity); 5. Coding bias: *variable* (except for the FISH dataset, which was set to *all*). Scripts to generate all NEXUS files and run analyses of individual data subsets were produced using the functions available in *ontobayes*.

## RESULTS

### Analyses of the BEE Dataset

Results from PROFILE analyses of the BEE dataset are shown in Table 1. Posterior coverage, i.e., the fraction of the total posterior probability distribution actually represented in the posterior sample of phylogenetic tree topologies (Lewis et al. 2016), for individual data subsets ranged from 51.0% for ‘mid leg’ to 99.7% for ‘male genitalia’; for profiles it ranged from 68.8% for ‘legs’ to 94.4% for ‘metasoma’. Such values indicate overall reasonable coverage (at least 50%) given that the number of possible phylogenetic tree topologies grows steeply with the increase in the number of taxa. BPI for individual data subsets ranged from 31.7% for ‘labrum’ to 80.7% for ‘male genitalia’ and for profiles from 39.6% for ‘legs’ to 62.1% for ‘metasoma’. Phylogenetic dissonance between different runs of MCMC for most individual data subsets was close to zero (0.1∼0.7%) indicating topological convergence in the posterior; the exceptions were ‘labrum’ and ‘mid leg’ showing slightly higher values (1.1% and 1.3% respectively). Dissonance for profiles ranged from 8.5% for ‘head’ to 14.7% for ‘mesosoma’ indicating substantial informational conflict among data subsets within profiles. A clustering dendrogram depicting hierarchical relationships among ontological terms annotated to individual data subsets included in PROFILE analyses is shown in Figure S1 (Supplementary Material: Fig. S1).

**TABLE 1.** Results from PROFILE analyses of the BEE dataset.

| Data subset | Coverage<sup>a</sup> | Information<sup>b</sup> | Dissonance<sup>c</sup> |
| --- | --- | --- | --- |
| **Mouthparts** |  |  |  |
| Labrum | 60.23 | 31.73 | 1.11 |
| Mandible | 94.39 | 60.70 | 0.38 |
| Maxilla | 96.29 | 58.69 | 0.27 |
| Labium | 97.45 | 65.60 | 0.14 |
| Sitophore | 89.16 | 55.67 | 0.55 |
| Run 1 | 85.26 | 47.60 | 13.61 |
| Run 2 | 85.31 | 47.49 | 13.83 |
| Mean | 85.28 | 47.54 | 13.72 |
| **Head** |  |  |  |
| Cranium | 97.42 | 69.55 | 0.17 |
| Tentorium | 82.71 | 47.00 | 0.63 |
| Run 1 | 87.34 | 54.64 | 8.39 |
| Run 2 | 86.86 | 54.57 | 8.62 |
| Mean | 87.10 | 54.61 | 8.51 |
| **Mesosoma** |  |  |  |
| Prothorax | 99.30 | 75.62 | 0.13 |
| Mesothorax | 97.81 | 67.56 | 0.22 |
| Metathorax | 85.32 | 49.73 | 0.66 |
| Run 1 | 91.93 | 58.25 | 14.61 |
| Run 2 | 91.96 | 58.39 | 14.78 |
| Mean | 91.94 | 58.32 | 14.70 |
| **Legs** |  |  |  |
| Fore leg | 83.36 | 50.07 | 0.49 |
| Mid leg | 51.00 | 30.25 | 1.28 |
| Hind leg | 95.01 | 59.14 | 0.33 |
| Run 1 | 69.23 | 39.61 | 12.04 |
| Run 2 | 68.34 | 39.67 | 12.05 |
| Mean | 68.78 | 39.64 | 12.05 |
| **Wings** |  |  |  |
| Fore wings | 75.31 | 40.77 | 0.55 |
| Hind wings | 91.23 | 55.19 | 0.37 |
| Run 1 | 71.45 | 42.24 | 10.37 |
| Run 2 | 71.29 | 42.23 | 10.38 |
| Mean | 71.37 | 42.23 | 10.37 |
| **Metasoma** |  |  |  |
| Female genitalia | 92.07 | 52.24 | 0.36 |
| Male genitalia | 99.70 | 80.85 | 0.14 |
| Run 1 | 94.44 | 62.19 | 11.84 |
| Run 2 | 94.31 | 62.09 | 11.94 |
| Mean | 94.37 | 62.14 | 11.89 |

<sup>a</sup> $\phi$ , estimated posterior coverage, expressed as percentage of maximum, as defined in Lewis et al. (2016).
<sup>b</sup> $BPI$ , estimated Bayesian phylogenetic information content, expressed as percentage of maximum, as defined in Lewis et al. (2016).
<sup>c</sup> $D$ , estimated phylogenetic dissonance, expressed as percentage of maximum, as defined in Lewis et al. (2016).

The dissonance dendrogram shows that data subsets included *a priori* in the same profile according to prior expert judgement about bee’s anatomy (Table 1) were not necessarily the ones less dissonant among themselves (Supplementary Material: Fig. S1). For example, subsets included in the ‘mouthparts’ profile (i.e., ‘labrum’, ‘mandible’, ‘maxilla’, ‘labium’, and ‘sitophore’) were not clustered in the dissonance dendrogram (Supplementary Material: Fig. S1: e.g., ‘labrum’ groups with ‘metathorax’ and ‘mandible’ with ‘fore leg’). This indicates that BPI content estimated from different subsets within profiles shows significant conflicting signal, i.e., information for alternative sets of phylogenetic tree topologies in the posterior distribution. Patterns observed in the dissonance dendrogram (Supplementary Material: Fig. S1) agreed with results shown in Table 1 indicating conflict among data subsets within profiles (phylogenetic dissonance >> 5%).

Results from ALL analyses of the BEE dataset (Supplementary Material: Fig. S2 and Table S1) showed similar patterns. Clustering of ontological terms annotated to data subsets based on phylogenetic dissonance does not reflect structural dependencies among anatomical entities of the bee anatomy. For example, BPI inferred from morphologically closely related entities such as ‘stipital sclerite’, ‘lacinial lobe’, and ‘galea’ (all *part_of* a bee ‘maxilla’) were highly dissonant among subsets (i.e., terms far apart in the dissonance dendrogram) whereas that of some unrelated entities such as ‘flabellum’ (*part_of* ‘labium’) and ‘female genitalia’ (*part_of* ‘metasoma’) were often less dissonant. BPI content inferred from individual subsets varied greatly in the ALL analyses of the BEE dataset as well (Supplementary Material: Table S1 and Fig. S2: barplots). Relative information, measured as the BPI of an individual subset divided by the mean BPI across all subsets, was particularly high for many subsets instantiating anatomical entities from the mouthparts (e.g., ‘sitophore’, ‘labrum’, ‘stipes’), prothorax (e.g., ‘profurcasternum’, ‘probasisternum’, ‘propleuron’), and metasoma (e.g., ‘male genitalia’, ‘female genitalia’) of bees (Supplementary Material: Fig. S2: bar heights higher than 1.0). BPI content for individual data subsets shown in Table S1 (Supplementary Material: Table S1) indicate considerably low phylogenetic information (< 25%) for at least half of them, also reflected in the higher phylogenetic dissonance values between different MCMC runs.

### Analyses of the FISH Dataset

As shown in Figures S3 and S4 (Supplementary Material: Figs S3 and S4), overall relationships among ontology terms were quite different between the semantic similarity and dissonance dendrograms indicating that phylogenetic information is not always structured by ontological knowledge and closely related terms in the ontology (i.e., semantically similar) do not always correspond to data subsets with more congruent phylogenetic information (i.e., lower phylogenetic dissonance). Relationships based on semantic similarity (Supplementary Material: Fig. S3), which reflect distances among concepts in the anatomy ontology, can be compared to relationships based on phylogenetic dissonance (Supplementary Material: Fig. S4), which reflect the degree of phylogenetic congruence or conflict among the posterior distributions of phylogenetic tree topologies obtained from the analyses of the subsets annotated to ontology terms. For example, the dissonance dendrogram indicate the following relationships among three particular anatomy terms: (‘premaxilla’ + ‘maxilla’) + ‘dentary’. This means that the posterior distributions of phylogenetic tree topologies obtained from the analyses of all characters annotated to the term ‘premaxilla’ and all characters annotated to the term ‘maxilla’ are more similar (i.e., include a more similar set of phylogenetic trees with similar posterior probabilities) than either is to the posterior distribution obtained from the analysis of all characters annotated to the term ‘dentary’. In other words, the phylogenetic information inferred from premaxillary and maxillary characters is more congruent; that for premaxillary and dentary or maxillary and dentary characters is less.

BPI content of individual data subsets and patterns of BPI and phylogenetic dissonance among-subsets mapped onto the semantic similarity dendrogram obtained for the FISH dataset varied greatly with most subsets presenting relatively low information (Fig. 4: middle column barplots). However, two major clusters of terms in the semantic similarity dendrogram (indicated by arrowheads) represent groups of relatively highly informative individual data subsets (e.g., some bones from the epibranchial and ceratobranchial series, maxilla, premaxilla, dentary, ectopterygoid and quadrate bones etc.). Relative information among-subsets, measured as among-subset BPI divided by mean among-subset BPI across all nodes of the semantic similarity dendrogram, was especially higher in some sectors of the dendrogram (Fig. 4a: blue circles) and decreased drastically towards deeper nodes (Fig. 4a: blue circles). Relative dissonance among-subsets, measured in a similar way, showed a similar but opposing pattern (as expected) with overall increase in values towards deeper nodes (Fig. 4b: red circles).

**FIGURE 4.**
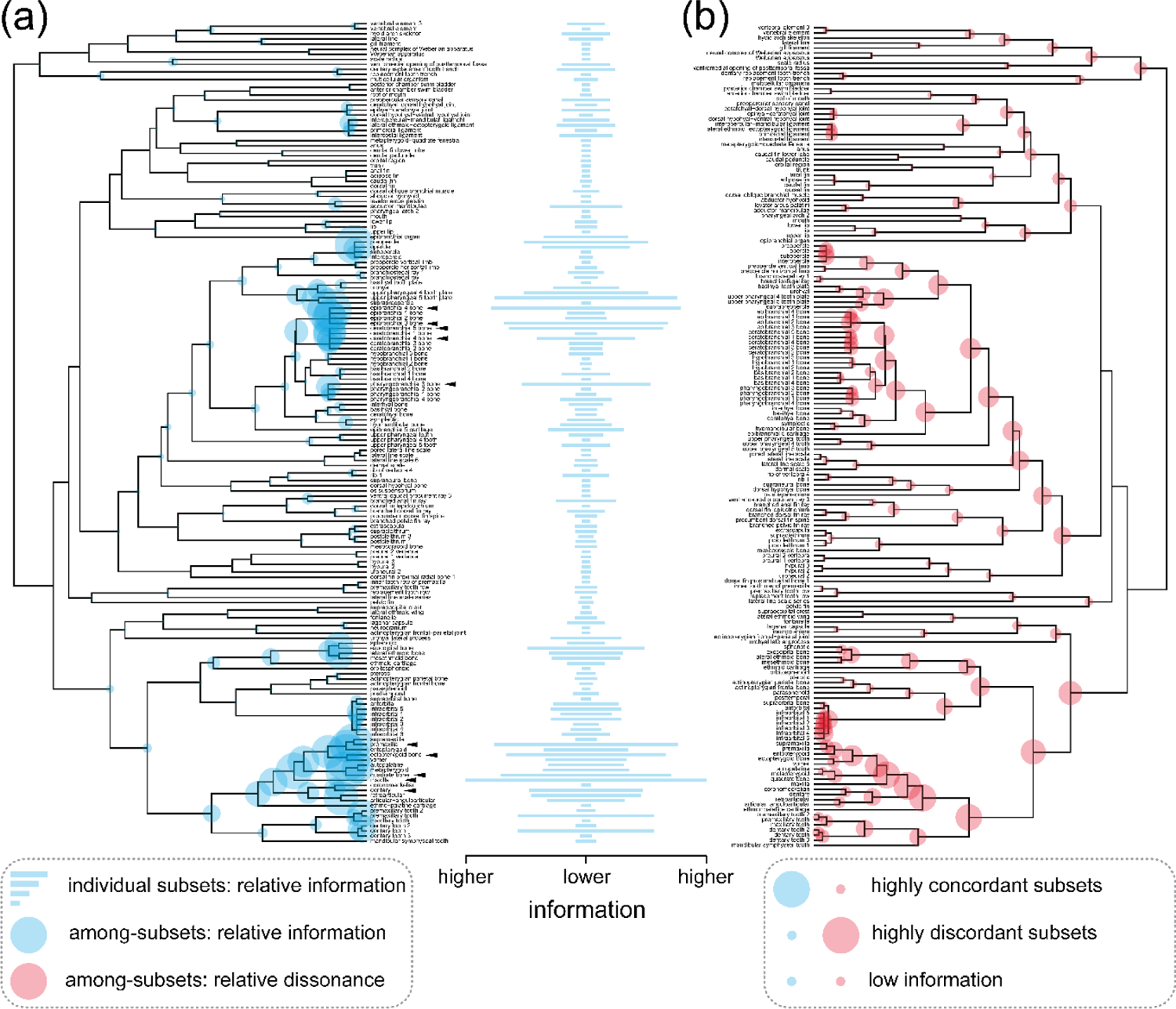
Bayesian phylogenetic information content for all anatomical entities linked to Uberon terms in the FISH dataset. Clustering dendrograms in (a) and (b) are obtained from pairwise semantic similarity between terms converted to a distance matrix. Barplots in middle column show information content of individual trait subsets defined by ontology terms relative to mean information across all subsets. Filled circles in trait dendrograms show (a) Bayesian phylogenetic information content and (b) phylogenetic dissonance among trait subsets defined by the ontology terms subtended by each node relative to respective mean values across all subsets. Bar lengths and circles have no absolute scale and are proportional to the relative maximum amount of (a) information or (b) dissonance observed. Bottom left and right boxes contain explanatory diagrams on how to interpret results in this figure. For colors, please refer to the online version of this paper available at XXX.

Patterns of among-subset information and dissonance are better understood in conjunction, as explained in the bottom-right box in Figure 4. Clusters of data subsets providing highly congruent phylogenetic information are also expected to present relatively higher among-subset relative information (Fig. 4a: large blue circles) and lower dissonance (Fig. 4b: small red circles) since they should represent similar posterior distributions of phylogenetic tree topologies (e.g., Fig. 2b: PCF and PVF). On the other hand, if data subsets provide highly conflicting information, then the opposite will be true, with relatively lower among-subset relative information (Fig. 4a: small blue circles) and higher dissonance (Fig. 4b: large red circles) (e.g., Fig. 2b: DF and IO). If most datasets provide little to no information at all, then both among-subset relative information and dissonance will be relatively lower since they should represent mostly flat, broadly overlapping, posterior distributions of phylogenetic trees. More complex scenarios, however, are usually found, with many clusters grouping multiple data subsets with varying degrees of information content and only partly overlapping posterior distributions of phylogenetic trees (e.g., Fig. 2b: PCF and DF) thus resulting in more ambiguous patterns of among-subsets relative information and dissonance, as observed for many nodes in Figure 4 (blue and red circles). Results were further inspected as phylogenetic tree topology trace plots (as available in the R package RWTY, Warren et al. 2017) to help assess degree of overlap between posterior distributions and better understand patterns of among-subsets information and dissonance. Some examples contrasting posterior distributions of phylogenetic tree topologies from both MCMC runs from the same data subset and from different subsets with congruent or conflicting phylogenetic information are provided in Figures S5 and S6, respectively (Supplementary Material: Figs S5 and S6).

Clade-specific phylogenetic information inferred from data subsets in the FISH dataset demonstrate that most phylogenetic information for the particular reference species tree obtained from the analysis of the full dataset (Fig. 5, bottom) is inferred from two major clusters of data subsets (Fig. 5, heatmap, dashed boxes) as indicated in the semantic similarity dendrogram (Fig. 5, right): one including bones from epibranchial, ceratobranchial, and pharyngobranchial series (Fig. 5, trait dendrogram, top cluster); and another including bones from maxilla, premaxilla, dentary, and infraorbital series, among others (Fig. 5, trait dendrogram, bottom cluster). With the exception of the first node (Fig. 5, species tree, bottom, N1), which was enforced due to rooting, only one node in the reference phylogenetic species tree received no support at all (Fig. 5, species tree, bottom, N7); all other nodes received variable amount of support from different subsets in both clusters (Fig. 5, top-left, heatmap; e.g., N2–N6; shade intensity proportional to posterior probability). The proportion of data subsets supporting each node in the phylogenetic species tree also varied (Fig. 5, bottom-right, barplots), with about only 1% of all subsets supporting N5 and between 4% and 9% supporting other nodes. It was also possible to investigate if the two inferred clusters of data subsets shared underlying ontological concepts. For example, we filtered all ontology terms defining subsets that are *part_of* ‘dermatocranium’ (Supplementary Material: Fig. S7: orange shaded rows in the heatmap) and found that information only from bones from the fish dermatocranium supported N5 (Supplementary Material: Fig. S7, species tree, bottom, N5) and most non-dermatocranium bones supported N2 and N4 (Supplementary Material: Fig. S7, species tree, bottom, N2 and N4).

**FIGURE 5.**
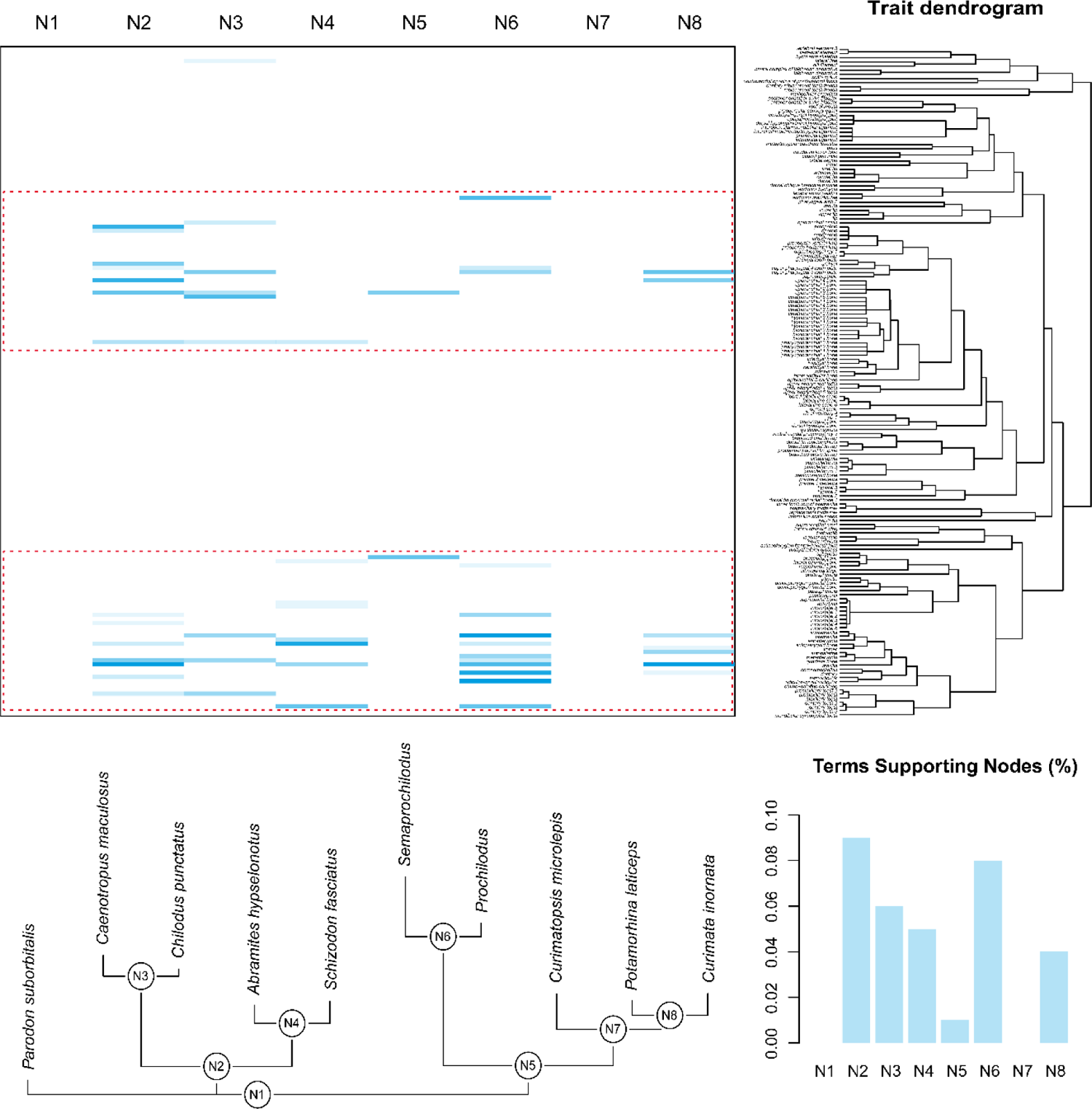
Clade-specific Bayesian phylogenetic information components in the FISH dataset. Heatmap shows which clades (columns) from a reference phylogenetic species tree (below) are supported by each subset defined by ontology terms (rows) in the reference trait dendrogram (right). Species tree is based on all characters. Trait clustering dendrogram is obtained from pairwise semantic similarity between terms converted to a distance matrix. Dashed boxes indicate two major clusters of data subsets. Heatmap color shade intensity is proportional to posterior probability. Barplots at bottom right show proportion of trait subsets supporting a given clade in the phylogenetic species tree. Abbreviations: N1…N8, nodes referring to clades in the phylogenetic species tree. For colors, please refer to the online version of this paper available at XXX.

### Analysis of the MINI Dataset

Results from the analysis of the MINI dataset (Supplementary Material: Fig. S8) showed little phylogenetic information to tree topology that could be useful to address the particular question about miniaturization in this sample of characiform fishes. Only a few data subsets (about 1%) provided information to N5, the clade enforcing the grouping of all miniature fishes (Supplementary Material: Fig. S8, species tree, bottom, N5). Most data subsets (about 7%) provided information to N2, the clade including all Characidae (Supplementary Material: Fig. S8, species tree, bottom, N2). No data subset provided information to N6 and N7 (Supplementary Material: Fig. S8, species tree, bottom, N6-N7), subclades of the miniature fishes clade. The majority of data subsets informative to nodes recovered in the reference phylogenetic species tree (Supplementary Material: Fig. S8, species tree, bottom, N2–N5) were annotated to ontology terms mostly related to anatomical entities comprising particular tooth rows from jaw bones (e.g., ‘premaxillary tooth row’, ‘maxillary tooth row’, ‘dentary tooth row’), with ‘premaxillary tooth row’ supporting the miniature fishes clade.

### Resampling Analyses

Resampling analyses show that mean values of BPI and phylogenetic dissonance were higher and lower, respectively, in datasets based on ontology term annotations compared to those composed by randomly resampling characters—strongly supporting that some ontology-based subsets carry shared phylogenetic information. BPI estimated for standard subsets were almost always higher than their respective resampled counterparts (Fig. 6a: e.g., PMX, MX, EB, CB). As expected, the opposite pattern was observed for phylogenetic dissonance, with most standard subsets showing lower values (Fig. 6b). A semantic similarity dendrogram for the selected ontology terms recovered two clusters, one for ‘premaxilla’ + ‘maxilla’ + ‘dentary’ + ‘infraorbital’ and another for ‘ceratobranchial bone’ + ‘epibranchial bone’ (Supplementary Material: Fig. S9). Results of pairwise comparisons between data subsets with estimates of phylogenetic dissonance obtained for standard and resampled subsets are shown in Figure S10 (Supplementary Material: Fig. S10). Note that estimates for comparisons between standard subsets show a trend of increasing phylogenetic dissonance (Supplementary Material: Fig. S10: e.g., PMX-MX, PMX-DEN, PMX-IO, PMX-EB etc.) when datasets annotated to increasing distantly related ontology terms were compared (Supplementary Material: Fig. S9).

**FIGURE 6.**
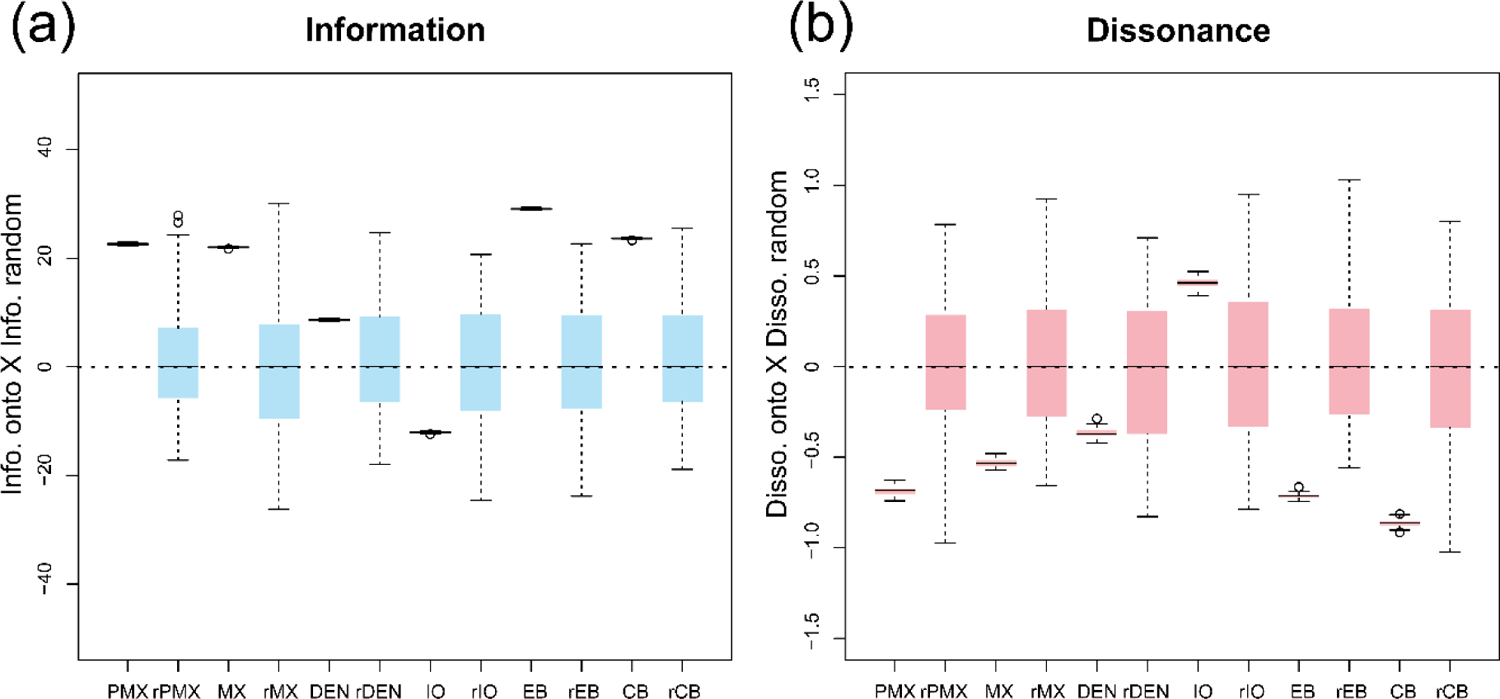
Boxplots showing estimated. (a) Bayesian phylogenetic information and (b) phylogenetic dissonance across replicated analyses for standard data subsets relative to resampled data subsets. Values above the dotted line indicate values higher than the median of the respective resampled data subsets. Note that information is higher and dissonance is lower for all ontology-based data subsets except IO than random subsets sampled of the same size, but without respect to ontology. Abbreviations: CB, ceratobranchial bone; DEN, dentary; EB, epibranchial bone; IO, infraorbital; PMX, premaxilla; MX, maxilla. The “r” prefix denote resampled subsets. For colors, please refer to the online version of this paper available at XXX.

## DISCUSSION

Ontologies bridge different domains of knowledge across life sciences (e.g., anatomy, development, genetics, behavior, ecology) allowing data integration within and across databases (Mabee et al. 2007; Deans et al. 2015). The recent growing interest in ontologies has contributed to the establishment of multiple collaborative projects targeting different biological entities (e.g., genes: The Gene Ontology Consortium 2000; cells: Bard et al. 2005; gross anatomy: Mungall et al. 2012), model organisms (e.g., mouse: Hayamizu et al. 2005; zebrafish: Sprague et al. 2008), and taxonomic groups (e.g., mammals: Smith et al. 2005).

Multispecies anatomy ontologies have been introduced for many taxa (e.g., amphibians: Maglia et al. 2007; fishes: Dahdul et al. 2010b; spiders: Ramírez and Michalik 2014; hymenopteran insects: Yoder et al. 2010) prompting assimilation of ontological knowledge in studies of evolutionary phenotypes (e.g., Mabee et al. 2012), semantic-aware anatomical descriptions (e.g., Mikó and Deans 2009; Silva and Feitosa 2019), and standardization of morphological terminology (e.g., Vogt 2008, 2009; Vogt et al. 2010; Karlsson and Ronquist 2012; Porto et al. 2016, 2017). In other words, ontologies are the structured knowledge that can be used to organize trait data in much the same way that phylogenies organize species data.

### Does ontology carry phylogenetic information?

In this work, we have asked to what degree phylogenetic information is structured by ontological knowledge by evaluating the BPI content and phylogenetic dissonance among ontology-annotated anatomical data subsets. If ontology carry any phylogenetic information, one would expect that sets of trees inferred from data subsets annotated with related ontology concepts would also be more similar (e.g., Fig. 2: PCF and PVF). In other words, their posterior distributions would concentrate probabilities in similar sets of trees. This would be indicated by high BPI and low phylogenetic dissonance among data subsets representing semantically similar concepts.

When analyzing the BEE dataset, through the PROFILE analyses, we find that subsets grouped based on anatomically related ontology concepts (i.e., ‘profiles’) actually exhibit considerably high phylogenetic dissonance (Table 1, values between 8.5∼14.7%). When analyzing the FISH dataset, we find substantial information in many ontology-annotated data subsets, but not universally across all anatomical subsets studied (Figs 4-5). Some clusters of similar ontology terms represent groups of highly informative individual data subsets (Fig.4: arrowheads) with high among-subset BPI (Fig. 4: blue circles) and moderate to low phylogenetic dissonance (Fig. 4: red circles). These clusters include concepts referring to some bones from jaws and branchial arches. These findings are consistent with the results from the resampling analyses of the FISH dataset, which show that BPI for data subsets containing characters annotated with the concepts of ‘maxilla’, ‘premaxilla’, ‘epibranchial’ and ‘ceratobranchial’ was higher than that of subsets based on a random resample of characters (Fig. 6b). The analyses of both datasets show that ontology does indeed structure phylogenetic information in some cases, thus prompting further investigation on the underlying biological processes that may explain that. However, ontology concepts and their relations do not fully explain phylogenetic information for all datasets and across all anatomical entities—as might be expected given the somewhat limited set of relations present in current anatomy ontologies. Instead, we observe that the semantic similarity dendrogram relating ontology concepts (Supplementary Material: Fig. S3) and the dissonance dendrogram relating posteriors inferred from the anatomical data subsets (Supplementary Material: Fig. S4) have very different topology. This indicates that additional processes or other biases are likely to also play a role in explaining BPI and dissonance values across anatomical subsets.

### How is phylogenetic information structured?

While we show that the ontology hierarchy does carry signal in the structuring of phylogenetic information for some datasets and anatomical concepts, it predictably does not do in all cases. Nevertheless, we can use the ontology hierarchy to interrogate morphological data with ontology knowledge in search for meaningful biological insights. Here, we asked if particular classes of anatomical entities were more phylogenetically informative than others.

As for the BEE dataset, for example, most information was inferred from anatomical entities instantiating concepts from mouthparts (e.g., ‘sitophore’, ‘labrum’, ‘stipes’), prothorax (e.g., ‘profurcasternum’, ‘probasisternum’, ‘propleuron’), and metasoma (e.g., ‘male genitalia’, ‘female genitalia’). As for the FISH dataset, two main clusters of anatomical entities (Fig. 5, heatmap, dashed boxes) provide most of the information for nodes recovered in the phylogenetic species tree (Fig. 5: bottom tree). One cluster includes many concepts from the jaw bones (e.g., ‘premaxilla’, ‘maxilla’, ‘dentary’); the other, many from the branchial arch bones (e.g., ‘pharyngobranchial’, ‘epibranchial’, ‘ceratobranchial series’); and most of these are developmentally associated with the dermatocranium (Supplementary Material: Fig. S7: orange shades). The two clusters of concepts and their association with ‘dermatocranium’ reinforce the findings that, for the FISH dataset, ontology seems to structure phylogenetic information. The analyses of both datasets show that indeed phylogenetic information is not uniformly distributed across anatomy ontology concepts.

Furthermore, anatomy entities do not provide the same information for all nodes in the phylogenetic species tree. For the FISH dataset (Fig. 5, bottom tree, N1-N8), for example, most information is inferred for N2-4 and N6, whereas N5 and N7 are inferred with little or no information from individual anatomy ontology concepts. This indicates that despite phylogenetic information not being uniform across all anatomical entities, it is still important to include a ‘semantic diversity’ of anatomical concepts in order to provide resolution for as many nodes as possible in the phylogenetic species tree.

### What sorts of processes may structure information?

If ontology hierarchy does not fully explain the phylogenetic information inferred from data, which other processes may explain it? Here we explored ontological knowledge summarized as a clustering dendrogram relating anatomical concepts by semantic similarity. This dendrogram was used as a proxy to describe anatomical/structural relations among real anatomical entities. These anatomical/structural relations might be interpreted as the product of developmental processes affecting morphogenesis of anatomical entities. Therefore, when we first asked the question whether ontology structures phylogenetic information, we were interested in knowing if anatomical/structural (∼developmental) relations among traits can influence their evolution. In other words, investigate if the evolution of some characters is non-independent due to anatomical/structural associations and/or other biological processes.

Non-independence among characters can result in more similar posterior distributions of trees inferred from dependent anatomical subsets—e.g., due to anatomical/structural associations. It may also result from common functional/ecological factors shared across species. Likewise, similar posterior distributions can simply be the result of shared evolutionary history. Some anatomical subsets may produce posterior distributions that are more congruent with the true species phylogeny (e.g., Fig. 1a,c). Others may agree within-subsets and/or among-subsets but disagree with the true species phylogeny (e.g., Fig. 1b,d). These can be easily accessed, for example, by contrasting posterior distributions for the species phylogeny—inferred from other sources of data (e.g., molecular data)—with posteriors inferred from data subsets annotated to each anatomical concept. Those agreeing with the species tree posterior distribution (i.e., high BPI and low dissonance) would indicate anatomical entities that evolved following the species phylogeny. Those disagreeing with the species phylogeny but agreeing among themselves (i.e., low BPI and high dissonance in relation to the assumed species tree posterior, but high BPI and low dissonance among themselves) would indicate anatomical entities that evolved under processes other than phylogeny, for example, concerted convergence due to shared functional/ecological factors across unrelated species (e.g., Fig. 1a-b, squares and circles). Then, for those subsets agreeing with the species phylogeny, it is possible to assess how much the phylogenetic information is structured by ontology by contrasting clustering dendrograms based on semantic similarity and phylogenetic dissonance (as discussed in previous sections). Finally, some anatomical concepts may be inferred with low information due to few characters in the respective data subsets and/or noise.

As it was shown before, the anatomical/structural ontology does indeed effectively cluster some groups of anatomical concepts by their patterns of phylogenetic information, but not for the entire anatomy. Conflict among anatomical subsets and the species phylogeny or shared response to convergent selective pressures are likely candidates to explain the evolution of these other traits. Indeed, results from the PROFILE analysis of the BEE dataset demonstrates the former scenario (Table 1). Posterior distributions inferred from anatomical entities associated with the same anatomy-based ‘profile’ (e.g., ‘mouthparts’, ‘head’, ‘legs’, ‘wings’ etc.) have high levels of dissonance with each other, indicating that BPI in this case is not structured by anatomical relations and there is considerable conflict among anatomical subsets. As for the FISH dataset, the two clusters of concepts (Fig. 5, dashed boxes) indicate that phylogenetic information is partly structured by ontology, as shown before, but also by the species history, since most anatomical subsets in such clusters are inferred with information supporting many nodes in the assumed species phylogeny (Fig. 5, bottom tree).

The MINI dataset shows an interesting case where the assumed species tree intentionally does not correspond to the most probable species phylogeny. By enforcing a clade grouping all miniatures (Supplementary Material: Fig. S8, N5), it was possible to observe different processes likely structuring the phylogenetic information of anatomical subsets. For example, a small cluster of related anatomical concepts referring to tooth rows from jaw bones of fishes (Supplementary Material: Fig. S8, dashed box) indicate some structuring of phylogenetic information by ontology, but not necessarily agreeing with a ‘true’ species phylogeny. On the other hand, several unrelated anatomical concepts provide phylogenetic information for Characidae (Supplementary Material: Fig. S8, species tree, bottom, N2), thus indicating congruence with the ‘true’ species phylogeny, but no *semantic signal* (i.e., ontology does not seem to structure phylogenetic information). Finally, characters from the anatomical concept ‘premaxillary tooth row’ support the miniature clade, thus indicating a possible case of concerted convergence due to miniaturization in such fishes.

### Alternative and complementary approaches

We acknowledge that some questions addressed here can be partially explored using existing or alternative methods. For example, there are different methods for assessing support to bipartitions (splits), compatibility and/or conflict among characters (Bandelt and Dress 1992: split decomposition; Hendy and Penny 1993: spectral analysis; Chen et al. 2005: spectral partitioning). These methods are not at odds with ours; they are complementary. Indeed we think they could also be enhanced by the inclusion of the ontology-guided approach. Furthermore, our analyzes are based on entropy-derived metrics of information and evaluate posterior distributions of tree topologies inferred from groups of characters (i.e., subsets), instead of character-by-character. This enables evaluation of how Bayesian (phylogenetic) information and conflict is structured by ontology and to make meaningful comparisons among data subsets.

Another important distinctive aspect of our approach is that it adopts the definition of “phylogenetic information” in the same sense as suggested in Lewis et al. (2016). Therefore, our approach assesses the (Bayesian) phylogenetic information of data subsets. This is useful because first, it considers not individual trees, but entire posterior samples, thus incorporate phylogenetic uncertainty; and second, it allows comparisons of how the information in different data subsets concentrate the probabilities from the prior set of possible tree topologies into a different (or similar) set of trees in the posterior. By guiding these comparisons with ontology knowledge and semantic distances, we can evaluate how independent *conceptual* modules support or disagree with each other and with the overall species tree topology—helping to alleviate a major challenge to morphological phylogenetics, the non-independence of characters.

### Limitations and caveats

One limitation of our approach is that it currently lacks a means to formally test for statistical significance of differences in BPI and dissonance values. Nonetheless, our intent was to help researchers assess the absolute and relative information/dissonance among ontology-annotated anatomical data subsets, and using ontologies to guide this exploration can help researchers to identify patterns across data subsets that might be explained by particular ontological relations and/or biological processes.

In our study, we used the Phenoscape Knowledgebase (KB: https://phenoscape.org) to calculate semantic similarity across all types of ontological relations present in the KB. We noted that semantic similarity values calculated did not always correspond to our *a priori* expectations in illuminating ways. For example, some characters annotated with different ontology terms may share high semantic similarity because they share *is_a* relationships with a particular ontology concept, such as characters annotated with terms that are subtypes of (i.e., *subclasses_of*) the concept ‘calcareous tooth’, despite being *part_of* anatomical structures in distinct body regions of a fish (e.g., ‘premaxillary tooth’, ‘maxillary tooth’, ‘dentary tooth’). This suggests that disentangling the different types of relations between terms (e.g., Vogt 2018a: *subsumption* vs. *parthood* relations) would allow for testing alternative hypotheses for the ontology structure and relations that best reflect the phylogenetic information inferred from anatomical data subsets. This would enable other types of hypotheses to be tested using phylogenetic character matrices.

Doing so would alleviate one potential critique of using semantic similarity dendrograms—the expectation that ontological relationships will fully describe the actual relationships among real anatomical entities instantiated by such terms. In fact, this should not be expected given that ontologies do not contain complete information, and because unlike phylogeny, there is no single bifurcating structure that can adequately describe all character relations. Furthermore, anatomical concepts available in an ontology can vary depending on the referential adopted (i.e., classification), terms can be characterized with varying degree of detail (i.e., granularity), and organismal anatomies can be represented in multiple alternative ways by different experts (i.e., semantic heterogeneity) (Vogt 2018b). Ontologies always reflects design decisions among its creators and maintainers and, therefore, there is no single correct scope or structure. For example, semantic similarity dendrograms, depending on the type of relations included in the reference anatomy ontology, may cluster terms such as ‘distal process of premaxilla’, ‘distal process of maxilla’, and ‘distal process of dentary’ because they all share the same *is_a* relationship (i.e., are different subtypes of) ‘distal process’ (i.e., *subsumption* relations), even though they are *part_of* different fish jaw bones (i.e., *parthood* relations). Nonetheless, potential biases due to ontology choice or character annotation with ontology terms can be directly assessed by comparing alternative ontologies in much the same way that alternative phylogenies (or phylogenetic networks) can be compared to assess how among-species variation is structured.

Another possible objection concerns the assumption that anatomical relationships always conform to hierarchies and, therefore, can be represented as dendrograms. Differently from species phylogenies, where the process of descent with modification produces a clear hierarchical pattern across species (for most organisms), for anatomical entities, such pattern may or may not be expected as a general rule for anatomical relationships. However, some studies do suggest this may be the case for some anatomical entities. For example, studies on the evolution of cell types and eyes in Metazoa show that relationships among some anatomical entities may in fact be well-represented as tree-like diagrams, both in developmental and evolutionary time (Oakley 2003; Arendt 2008; Arendt et al. 2016). Nevertheless, much like genetic data with frequent horizontal gene transfer, *semantic signal* will often likely require multiple topological structures to best explain and predict character similarity. We argue that interrogating datasets with these alternative sets of relations and topologies is likely to reveal much about the processes governing morphological evolution, and argue for the continued development of robust ontologies for organismal traits.

### Perspectives and future directions

Applying ontology-guided approaches and moving beyond the flat, one-dimensional partitioning of characters has enormous potential for making sense of trait evolutionary patterns. For example, one can assess the phylogenetic information provided by data subsets annotated to particular ontology terms in respect to one or more nodes of interest in a given reference phylogenetic species tree (e.g., Fig. 5, bottom, species tree). Node(s) in such trees may characterize clade(s) of organisms sharing a particular biology or some traits of relevance; and by interrogating this node, we can discover and identify subsets of morphological characters that are phylogenetically highly informative for that particular node (e.g., Supplemental Material: Fig. S8: MINI dataset). Such an approach can be expanded and generalized to any test statistic of interest that can be calculated across the phylogeny or on a per character basis. For example, a researcher might be interested in the magnitude of support for a rate shift at a particular node, rather than the BPI content at the node given a particular reference ontology.

Such metrics can then be evaluated in light of the relationships among terms annotated to character data subsets, including using different ontological relations (e.g., *part_of*, *develops_from*) or distance metrics (e.g., *Jaccard*, *Resnik*) to build a semantic similarity dendrogram. This can mirror the way that alternative phylogenetic tree topologies are used to assess and compare phylogenetic information and signal across species, and they can shed light on the underlying processes determining similar evolutionary patterns in morphological traits. This approach can be employed, for example, to investigate if highly (or alternatively slightly) informative data subsets annotated with particular anatomical terms share any common underlying ontological relations. For example, we observed that most characters informative for the FISH dataset are included in data subsets defined by ontology concepts referring to bones that are *part_of* ‘dermatocranium’(Supplemental Material: Fig. S7) thus indicating possible structural/developmental dependencies among such traits.

Future research on Bayesian phylogenetic information will likely help to circumvent the limitation to small datasets by using tree priors allowing for polytomies or better strategies to sample posterior probability distributions (see discussions in Lewis et al. 2016). Further studies could make use of alternative visualization graphs for the relationships among ontology terms, using networks instead of dendrograms, and the selection of specific types of ontological relations, distance metrics, or subgraphs to represent the ontology structure (see also Vogt 2018b for additional insights into using graphs in ontology-aware phylogenetic analysis).

## SUPPLEMENTARY MATERIAL

Data available from the Dryad Digital Repository: http://dx.doi.org/XX.XX/XXX.[XXXX].

## ACKNOWLEDGEMENTS

We would like to thank the editors, Dr. Lars Vogt, and an anonymous reviewer for the thoughtful revisions on this manuscript. We are thankful to members of Uyeda’s lab for valuable comments and suggestions on early drafts. DSP would like to specially thank Matthew W. Pennell and his lab for the insightful discussions in their lab meeting. DSP is grateful to Fabio A. Machado for helping fixing issues with the code used in *ontobayes*.

